# Label-Free Optical Mapping for Large-Area Biomechanical Dynamics of Multicellular Systems

**DOI:** 10.1101/2024.06.12.598186

**Authors:** Yen-Ju Lin, Xing Haw Marvin Tan, Yijie Wang, Pei-Shan Chung, Xiang Zhang, Ting-Hsiang Wu, Tung-Yu Wu, Arjun Deb, Pei-Yu Chiou

## Abstract

Mapping cellular activities over large areas is crucial for understanding the collective behaviors of multicellular systems. Biomechanical properties, such as cellular traction force, serve as critical regulators of physiological states and molecular configurations. However, existing technologies for mapping large-area biomechanical dynamics are limited by the small field of view and scanning nature. To address this, we propose a novel platform that utilizes a vast number of optical diffractive elements for mapping large-area biomechanical dynamics. This platform achieves a field-of-view of 10.6 mm X 10.6 mm, a three-orders-of-magnitude improvement over traditional traction force microscopy. Transient mechanical waves generated by monolayer neonatal rat ventricular myocytes were captured with high spatiotemporal resolution (130 fps and 20 µm for temporal and spatial resolution, respectively). Furthermore, its label-free nature allows for long-term observations extended to a week, with minimal disruption of cellular functions. Finally, simultaneous measurements of calcium ions concentrations and biomechanical dynamics are demonstrated.

## Introduction

To maintain vital functions, a multicellular system requires coordinated interplays among millions to billions of cells to sense, adapt, and probe their surrounding microenvironments^1^. Conduction velocities among cells can range from sub-meters to a few meters per second for cells such as nerve fibers and cardiomyocytes^2,3^. Abnormal conduction blocks or un-orchestrated conduction activities of excitable cells in localized areas can lead to fatal failure of the entire multicellular system^4,5^. Therefore, large-area mapping of cellular activities with high spatiotemporal resolution is essential to identify localized heterogeneities and their corresponding contributions to the collective behaviors exhibited by the multicellular system.

Dynamic interaction and progression of cellular activities can be revealed by measuring various bio-indictors, such as the biomechanical, biochemical, and bioelectrical properties of cells. For example, large-area dynamic changes of bioelectrical and biochemical properties like transmembrane voltage or ion concentrations are often measured by conventional optical mapping using fluorescent dyes or microelectrode arrays^6,7^. While these techniques are well-established, the tools to measure large-area biomechanical properties, an equally crucial indictor of cellular activities, are still lagging a lot behind. Biomechanical properties, such as cellular stiffness and cell-generated force, play pivotal roles in mechanotransduction, the process cells sense, adapt and respond to external stimuli in their surrounding microenvironments^8^. They are critical measures that can indicate the physiological states and molecular configurations of cells^9,10^. However, existing technologies to measure the dynamics of biomechanical properties are limited to either a small number of cells or the bulk properties of cells without detailed spatial information^11,12^. For instance, traction force microscope (TFM), the classical technique to measure cellular traction force, typically has field-of-view smaller than 1mm^2^ and can handle less than 50 cells such as fibroblasts and smooth muscle cells per experiment^13–15^. Recently, several optics-based approaches have emerged, incorporating the utilization of Moiré patterns and diffractive gratings^16,17^. Nevertheless, since the contributions from individual cells to the resultant optical interference are not decoupled, these methods are either suitable for studying a limited number of cells or exhibit poor spatial resolution. On the other hand, as an alternative approach, calcium ion (Ca^2+^) concentration is usually measured as an indirect indication of biomechanical properties. This is because Ca^2+^ is an essential precursor for the contraction of muscles cells. However, there have been studies suggesting that calcium concentration is a limited predictor for quantifying cellular traction forces^18^. Hence, direct measurement of biomechanical properties is still desired to study the physiological interaction of calcium ion and the subsequent mechanical activities.

Motivated by the necessity to directly profile biomechanical dynamics across a large area, a platform called Single-Pixel Optical Tracers (SPOT) has been proposed recently^19^. Leveraging the independent operation of numerous grating disks, this platform eliminates the tedious steps to pinpoint the location of each sensing unit. Instead, it translates the traction force generated by cells into optical color spectra for comprehensive large-area profiling. Notably, cellular force measurements have been successfully demonstrated with a field-of-view over 13mm^2^ at 83 frames per second while maintaining a sub-cellular resolution. Nonetheless, the chromatic dispersion stemming from a wide spectrum illumination source ultimately limits its ability to further enhance the throughput and be compatible with other common microscopes. Here, to address these limitations, we propose a new platform that employs optical diffraction patterns instead of color spectra. This platform utilizes a massive number of optical diffractive elements embedded periodically in an elastic membrane to track the traction force generated by the cells seeded on top. To demonstrate the capabilities of our platform, monolayer tissue comprising a sub-million number of neonatal rat ventricular myocytes (NRVMs) is seeded onto our devices. An observation field of view over 110 mm^2^ is achieved, which is 3 orders of magnitude improvement compared to traditional TFM. Meanwhile, high spatiotemporal resolution is maintained, achieving a frame rate of 130 frames per second and a spatial resolution of 20 µm. This facilitates concurrent recording of collectively beating spanning over large area and the identification of local heterogeneities in mechanical waves propagation. Moreover, featuring as a label-free approach, this platform is capable of monitoring samples under native conditions or conducting various treatments over extended durations. Finally, we integrated the platform with a fluorescent imaging system to simultaneously track the transit of calcium ion concentrations along with the biomechanical dynamics from the beating of NRVMs. This represents the first demonstration of detailed mechanical wave propagation over a long period of time and the corresponding calcium ion transient. The results emphasize the potential of our platform for applications such as drug screening and pathological studies.

## Results

### Large-Area Mapping of Biomechanical Dynamics via Array of Diffractive Elements

A conceptual schematic of the proposed platform to conduct large-area mapping of biomechanical dynamics is illustrated in **Fig. 1**. The platform employs a multitude of optical diffractive elements embedded periodically in an elastic membrane (polydimethylsiloxane (PDMS)) to monitor the traction forces exerted by the cells seeded on top. A collimated light beam illuminates the diffractive elements, and the resulting diffraction patterns of each element are detected by the image sensor below. The diffractive elements are responsible for translating the lateral displacements induced by cell-generated forces to discernible changes that can be measured by the image sensor. As depicted in **Fig. 1b**, each diffractive element consists of a circular metal disk on the top layer of PDMS and a metal sheet with a central circular aperture at the bottom. In other words, the top and bottom metal layers form complementary shapes in terms of its opacity but are vertically offset by the PDMS membrane. Since the bottom metal layer is on the glass substrate, its movement is restrained. In the absence of applied force, most of the illuminated light is blocked, with only a small portion of diffracted light penetrating through the edge. When cells apply traction force on the PDMS membrane, the top metal layer moves while the bottom metal layer remains fixed. This movement creates eclipse-like apertures of varying sizes and generates distinct diffraction patterns for each diffraction element (**Fig. 1c**). Consequently, the transmitted light intensity and the diffraction pattern of each element serve as direct indicators of the local cellular traction force applied to the PDMS membrane.

**Fig. 1.**
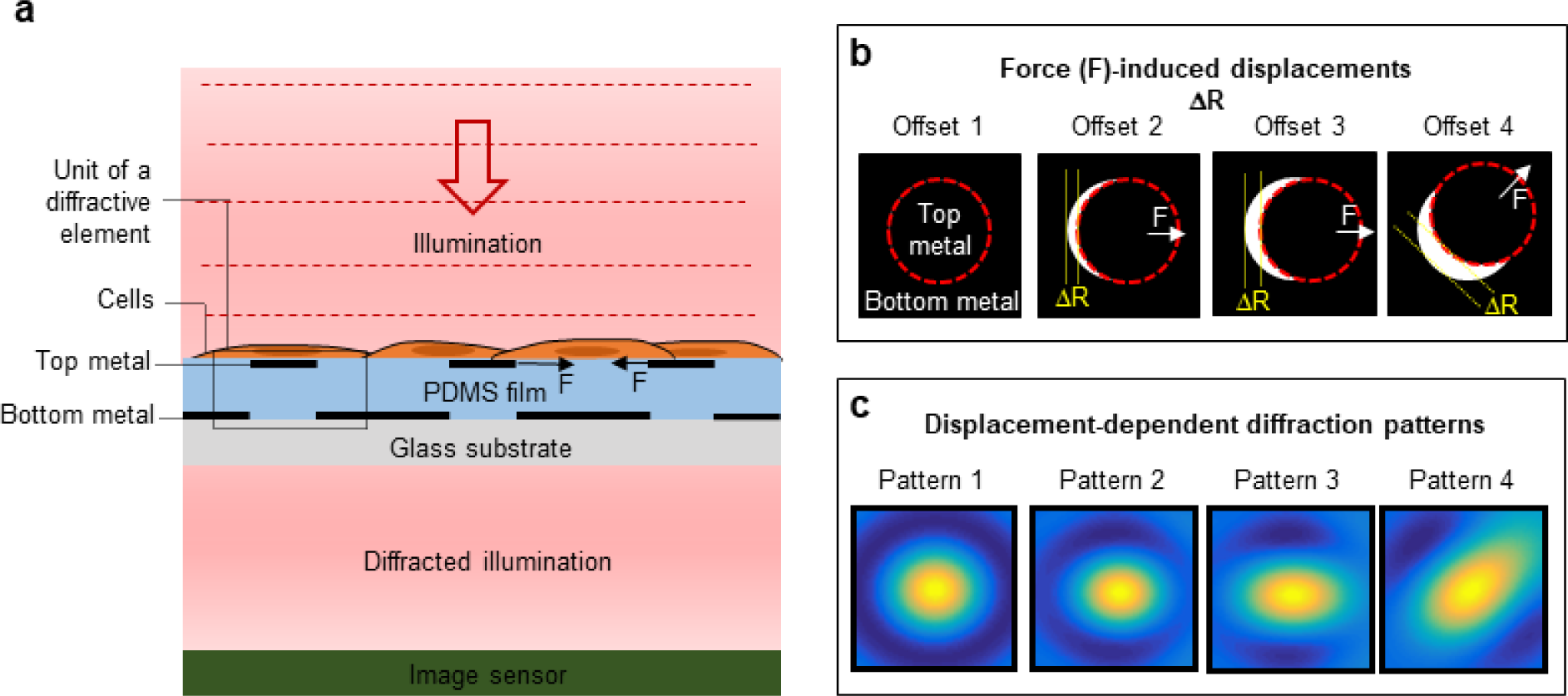
Schematic and working principle of the label-free optical mapping. **a.** Illustration of the system for label-free optical mapping of large-area biomechanical dynamics. A uniform collimated illumination with wavelength of 660nm is incident onto the monolayer cell sheet seeded on the PDMS film with embedded diffractive elements. Cellular traction forces induce lateral displacements ΔR on the diffractive elements. These displacements result in different diffraction patterns, which are recorded by the image sensor below. This translation from displacement to optical properties bypasses the limitations of pinpointing minute displacements and allows for parallel processing of numerous diffractive elements, leading to a significant enhancement of the observation field-of-view **b.** and **c.** Four representative examples of force-induced displacements and the displacement-dependent diffraction patterns, respectively. The intensity and shape of the diffraction pattern are directly related to the magnitude and direction of the cell-generated force.

### Real-time visualization of mechanical wave propagation over large area with high spatial resolution

To exhibit the capability of our platform in capturing rapidly propagating and transient mechanical waves among large cell populations, we chose to measure the cardiac beatings resulting from monolayers of NRVMs (refer to **Methods** for cell-seeding details). **Fig. 2a** displays three consecutive cardiac beatings extracted from a diffractive element stemming from a NRVM monolayer. The zoom-in figure illustrates the definition of activation time and activation duration, which are critical for evaluating the spatial evolution and homogeneity of cardiac beating. Activation time characterizes the timing when the beating wave arrives at a specific position, while activation duration indicates how long the beating lasts at a given location. Both parameters are functions of spatial coordinates. It should be noted that we employed activation time during the upstroke phase to demonstrate our platform’s capability in studying the propagation of mechanical waves. Alternatively, researchers can explore other crucial time points, such as the downstroke time and repolarization midpoint, to investigate various physiological stages during cardiac beats ^20,21^.

**Fig. 2.**
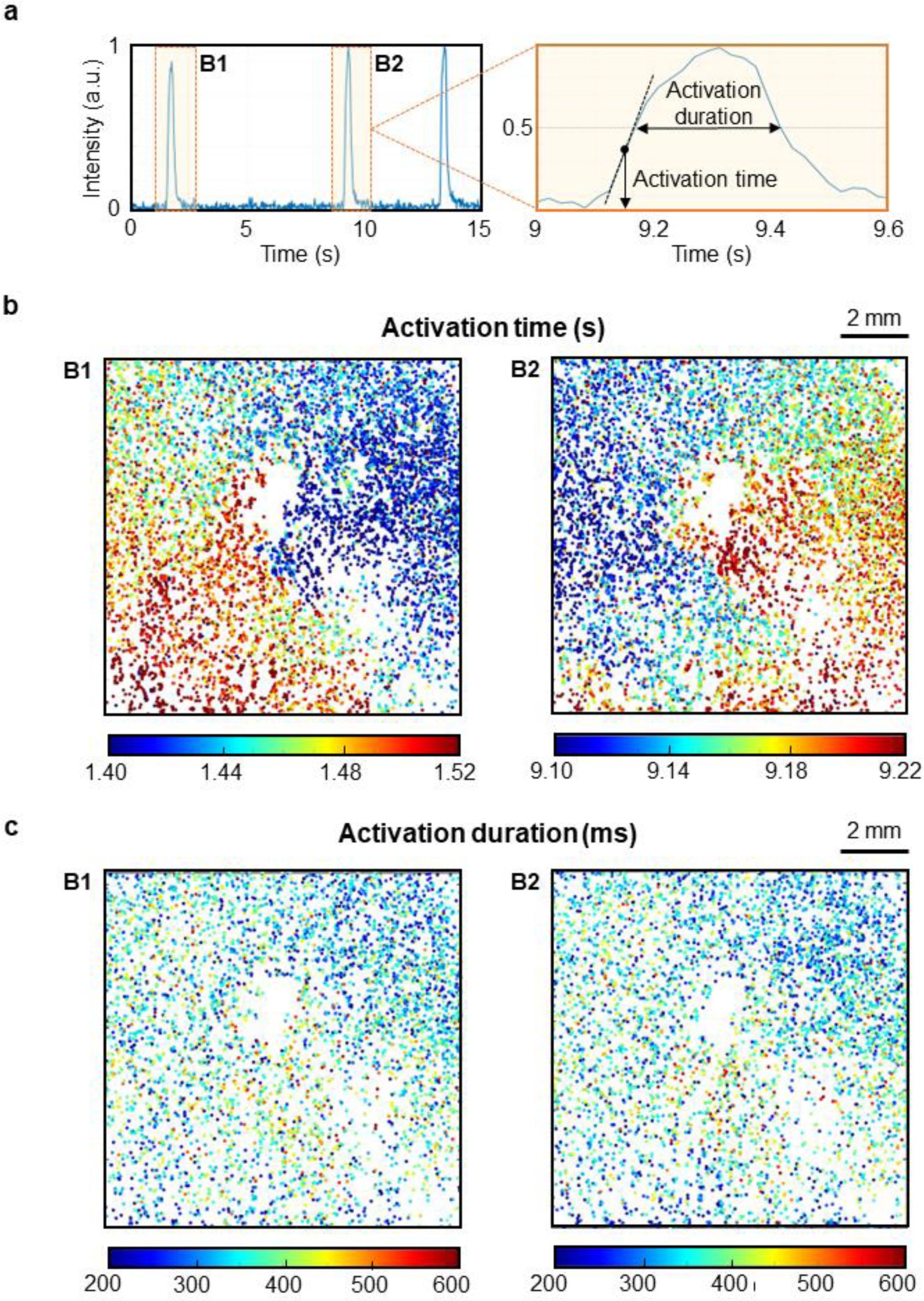
Real-time visualization of mechanical wave propagation over large area with high spatial resolution. **a.** Three consecutive cardiac beatings extracted from a diffractive element stemming from a NRVM monolayer. The zoom-in figure illustrates the definition of activation time and activation duration, which are critical for evaluating the spatial evolution and homogeneity of cardiac beating. Activation time characterizes the timing of the beating wave’s arrival at a specific position, while activation duration indicates the duration of the beating at a given location. **b.** Activation maps of the first two consecutive beats. Despite occurring sequentially in time, they exhibit completely distinct spatial patterns. This transient beat-to-beat variation emphasize the necessity for tools capable of measuring the large-area phenomena in a single shot without scanning. **c.** Activation duration maps of the first two beats. While activation time maps displays significant variation, activation duration maps for these two beats show minimal differences.

**Fig. 2b** shows the activation maps of the first two beatings of **Fig. 2a**. Although they occur sequentially in time, they feature completely distinct spatial patterns. The first beating is initiated roughly from the center and propagates along both the right and left arc to the bottom left end, while the onset of the second and third beating is from the left-hand side, with the wave traveling to the right-hand side. The videos displaying the time-course intensity changes of the first and second beatings are provided in **Supplementary Movie S1**. In addition to the global trends of beating patterns, we can observe numerous heterogenous points that do not follow the main beating from the bulk, thanks to the high spatial resolution of our platform. **Supplementary Figure S2** presents several represented heterogeneous traces, revealing the diverse contours of intensity changes over time, including the variations in the number of beatings, trace shape and signal to noise ratio. These variations are clear indications that the strength and profile of cellular traction force vary considerably among cell populations. Therefore, high spatial resolution is essential for large-area techniques to study the status of individual cells. On the other hand, while there is a significant variation in activation time maps, activation duration maps of these two beatings (given in **Fig. 2c**) do not exhibit significant differences.

### Continuous monitoring of mechanical dynamics during cardioversion via electrical stimulation

The spiral wave of cardio beating is often associated with abnormal cardiac activities or arrhythmia ^22,23^. Our platform has captured this spiral pattern and subsequently observed cardioversion after applying electrical stimulation. The stimulation schematic is displayed in **Fig. 3a**. The stimulating electrodes are positioned approximately 2mm away from the active area on the right. A 1Hz, 50% duty cycle square wave with a peak-to-peak voltage of 200 mV is utilized to pace the sample for 1 min. **Fig. 3b** demonstrates the time-course intensity distributions of the sample before and after the electrical stimulation (Videos are given in **Supplementary Movie S2**). Prior to stimulation, the spiral wave initiated slightly to the left of the center and rotates counterclockwise. Following the electrical stimulation, the waveform starts to propagate from right to left, corresponding to location of the stimulating electrode on the right-hand side of the sample. These results strongly demonstrate our system’s capabilities to track large-area biomechanical dynamics in real time.

**Fig. 3.**
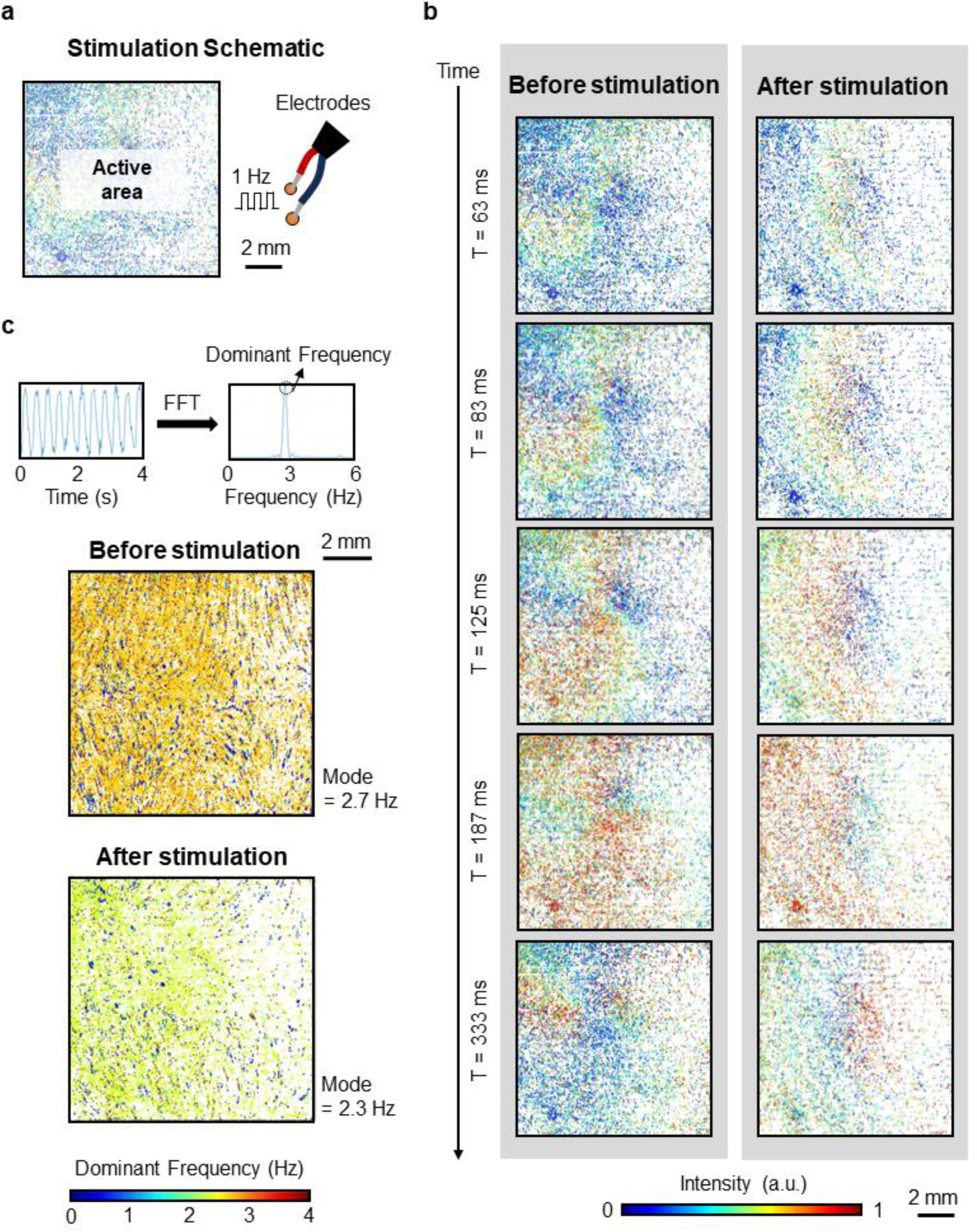
Continuous monitoring of mechanical dynamics during cardioversion via electrical stimulation. **a.** The stimulation schematic. The stimulating electrodes are positioned approximately 2mm away from the active area on the right. A 1Hz, 50% duty cycle square wave with a peak-to-peak voltage of 200 mV is utilized to pace the sample for 1 min. **b.** The time-course intensity distributions of the sample before and after the electrical stimulation. Prior to stimulation, the spiral wave initiated slightly to the left of the center and rotated counterclockwise. Following the electrical stimulation, the waveform started to propagate from right to left, corresponding to the location of the stimulating electrode on the right-hand side of the sample. These results strongly demonstrate our system’s capabilities to track large-area biomechanical dynamics in real time. **c**. Definition of dominant frequency calculation and the dominant frequency maps before and after electrical stimulation.

Next, the dominant frequency of the constantly beating cells is calculated before and after electrical stimulation. This is accomplished by performing fast Fourier transform (FFT) of the time-course intensity traces from each diffractive element. The frequency component with the largest intensity value is considered the dominant frequency for that element. The dominant frequency maps before and after electrical stimulation are given in **Fig. 3c**. The mode values (most frequent number) are 2.7 Hz and 2.3 Hz, respectively. Our platform’s high spatial resolution and large area features enable us to discern both the main beating frequency exhibited by the majority of cells and various heterogeneous behavior, which may express as slower or faster beats compared to the majority. This will be challenging to achieve for both traditional traction force microscopes or conventional optical mapping techniques using fluorescent dyes, in which the spatial resolution and large area cannot be attained simultaneously.

### Long-term observation of biomechanical dynamics under native conditions

As cell cultures grow and mature over time, recording their biomechanical dynamics without disrupting their physiological activities can effectively eliminate uncertainties associated with external factors such as the impacts of fluorescence dyes on cells. This enables a more straightforward recognition of the causality of biological events and pathological progression. Hence, in this subsection, we highlight our platform’s strength in conducting long-term observation of samples under native conditions by continually logging the biomechanical dynamics of the same cell culture daily for up to 7 days, which cannot be achieved by conventional optical mapping using fluorescent dyes. Limitations such as dye internalization, photobleaching and phototoxicity typically restrict the maximum observation duration to only several hours ^7,24^.

In the following discussions, the day when the NRVM cells were isolated and seeded on the device is denoted as Day 0. The cells usually begin beating within 24 to 48 hours after seeding. Our measurements started on Day 3 and were repeated daily until Day 9. On Day 7, we conducted a chemical stimulation using a low concentration of phenylephrine and recorded the corresponding response. Phenylephrine, being a kind of α-1 adrenergic receptor agonist, frequently induces hypertrophic responses in cardiomyocytes^25^. **Fig. 4a** displays the maps of activation time, each accompanied by a representative time-course intensity trace for each day. There is a defective region in the top left corner of this device, rendering the data (diffraction patterns) inaccessible. However, the cells are present in that area. **Fig. 4b** presents the dominant frequency for each day with a zoom-in figure illustrating the change associated with the chemical stimulation performed on Day 7.

**Fig. 4.**
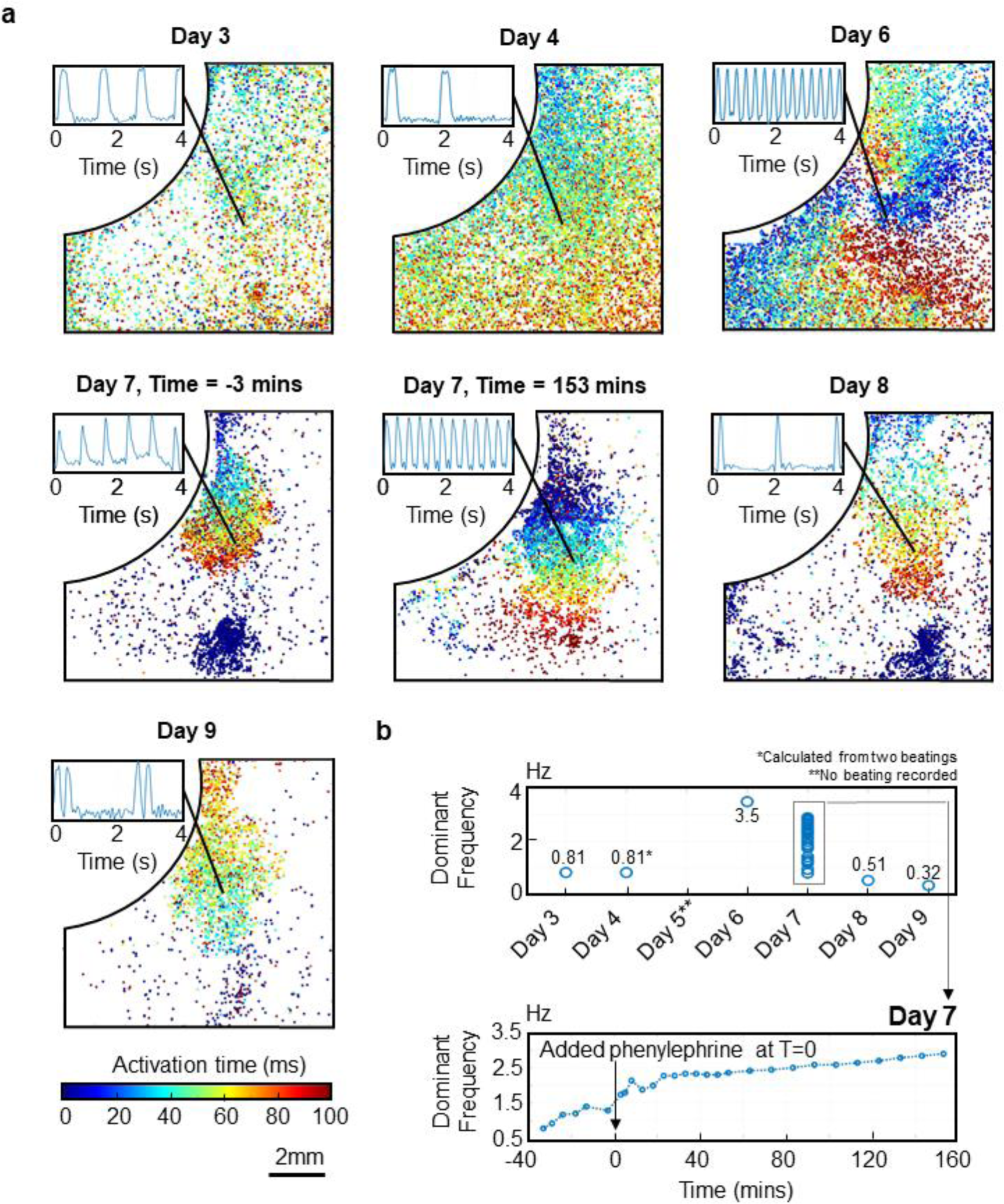
Long-term observation of biomechanical dynamics under native conditions. **a.** The maps of activation time, each accompanied by a representative time-course intensity trace in the inset for that day. A defective region in the top-left corner of this device renders the diffraction patterns inaccessible, although the cells were present in that area. Day 3 and Day 4 present similar spatial patterns of wave propagation. While constant beatings were observed on Day 3, irregular beating was recorded on Day4. No clear beatings were observed on Day 5. On Day 6, a spiral wave pattern with a beating rate of 4.4 Hz appeared. On Day 7, a low-concentration phenylephrine was added to the sample as chemical stimulation. Just before the drug administration (time = -3 minutes), two clusters were observed at the top right and top bottom, beating with a rate of 1.29 Hz. By the end of the treatment, the beating frequency increased to 2.89 Hz, with the two clusters becoming more connected. The drug was then removed, and fresh medium was replenished to the cells culture. After overnight incubation, the recording resumed on Day 8. At this time, the connected clusters separated, with a longer delay between their activation time. Lastly, on Day 9, only the cluster in the top-right corner remained constantly beating at 2.5Hz. **b.** The dominant frequency for each day, with a zoom-in figure illustrating the change associated with the chemical stimulation performed on Day 7.

As can be seen in **Fig. 4a**, on Day 3, regular beatings with a rate of 0.81 Hz were observed. The onset of the beating was from the top-left and propagated to the bottom-right corner. Then, on Day 4, the beating pattern followed the same spatial arrangement, traveling from the top-left to the bottom-right. However, the beating of NRVM is not consistent in culture, with only two beats occurring over the entire 16.8-second recording. Although the beating activity was not steady, the captured two beatings had similar time difference as those on the previous day (1.25s). Interestingly, the activation time map shows denser points compared to that of Day 3. This may indicate that more cells were engaged in the beating activities, resulting in large traction forces being applied on the device to activate more diffractive elements. Next, no clear beating of the cells was recorded on Day 5. Subsequently, on Day 6, a spiral pattern appeared with a beating rate of 4.4 Hz. Afterward, chemical stimulation was performed on Day 7. Specifically, a low-concentration (1 µM) phenylephrine (PE) was added to the culture medium and treated for 153 minutes. Just before the drug administration (time = -3 minutes), two clusters were observed at the top-center and bottom-center, beating with a rate of 1.29 Hz. Phenylephrine is an alpha agonist and known to increase the frequency of beating cardiac muscle cells in culture. By the end of the treatment, the beating frequency increased to 2.89 Hz, with the two clusters becoming more connected. The drug was then removed, and fresh medium was replenished to the cell culture. After overnight incubation, the recording resumed on Day 8. At this time, the connected clusters separated, with a longer delay between their activation time. The majority of cells beat at 0.51 Hz. Lastly, on Day 9, only the cluster in the top-center remained constantly beating at 2.5Hz. The beating was initiated from the bottom of the cluster and traveled to the top, opposite to the directions shown in the previous few days. Additionally, it is worth noting that there were two consecutive peaks associated with each beating.

In summary, long-term observations of large-area biomechanical dynamics are achieved with the span extended to a week, with minimal disruption to the physiological activities of cells. It is crucial to emphasize that the designed experiment is not aimed at drawing any biological conclusions from a single sample, but rather to demonstrate our platform’s capabilities to dynamically monitor cells under native conditions over an extended period. By continuously observing biomechanical dynamics and minimizing disturbances to cell cultures, our platform holds great potential for applications in studying disease development and evaluating the effectiveness of drugs.

### Simultaneous measurements of calcium ions concentrations and biomechanical dynamics

Simultaneous measurements of multiple parameters, encompassing bioelectrical, biomechanical, and biochemical properties, empower us to investigate the cellular system from diverse perspectives and understand the interplays between these properties. For instance, transmembrane voltage and calcium ion concentration are commonly reported together in the literature^7,24^. However, concurrent mapping of biomechanical dynamics with other parameters over a large area is rarely explored. Consequently, we expanded our platform to integrate the optics required for fluorescent imaging. As a proof-of-concept, we conducted simultaneous measurements of calcium ion (Ca^2+^) concentration and biomechanical dynamics. The high-affinity calcium ion dye, Rhod-2, was utilized for fluorescent imaging (Refer to **Methods** for the dye loading protocol and setup for fluorescent imaging).

**Fig. 5a** displays the time-course intensity distributions, revealing the biomechanical dynamics measured by our platform using diffractive elements. A spiral wave is clearly visible across the entire field of view. In **Fig. 5b**, the time-course intensities indicating Ca^2+^ concentration (Orange curves, labelled as “F[Ca^2+^]) and biomechanical dynamics (blue curves) from various locations are compared. It is evident that the fluorescent signal and the biomechanical signal exhibit similar profiles, including certain elongated periods. This observation verifies that calcium concentrations and biomechanical dynamics are closely related in the physiological activities of muscle contractions.

**Fig. 5.**
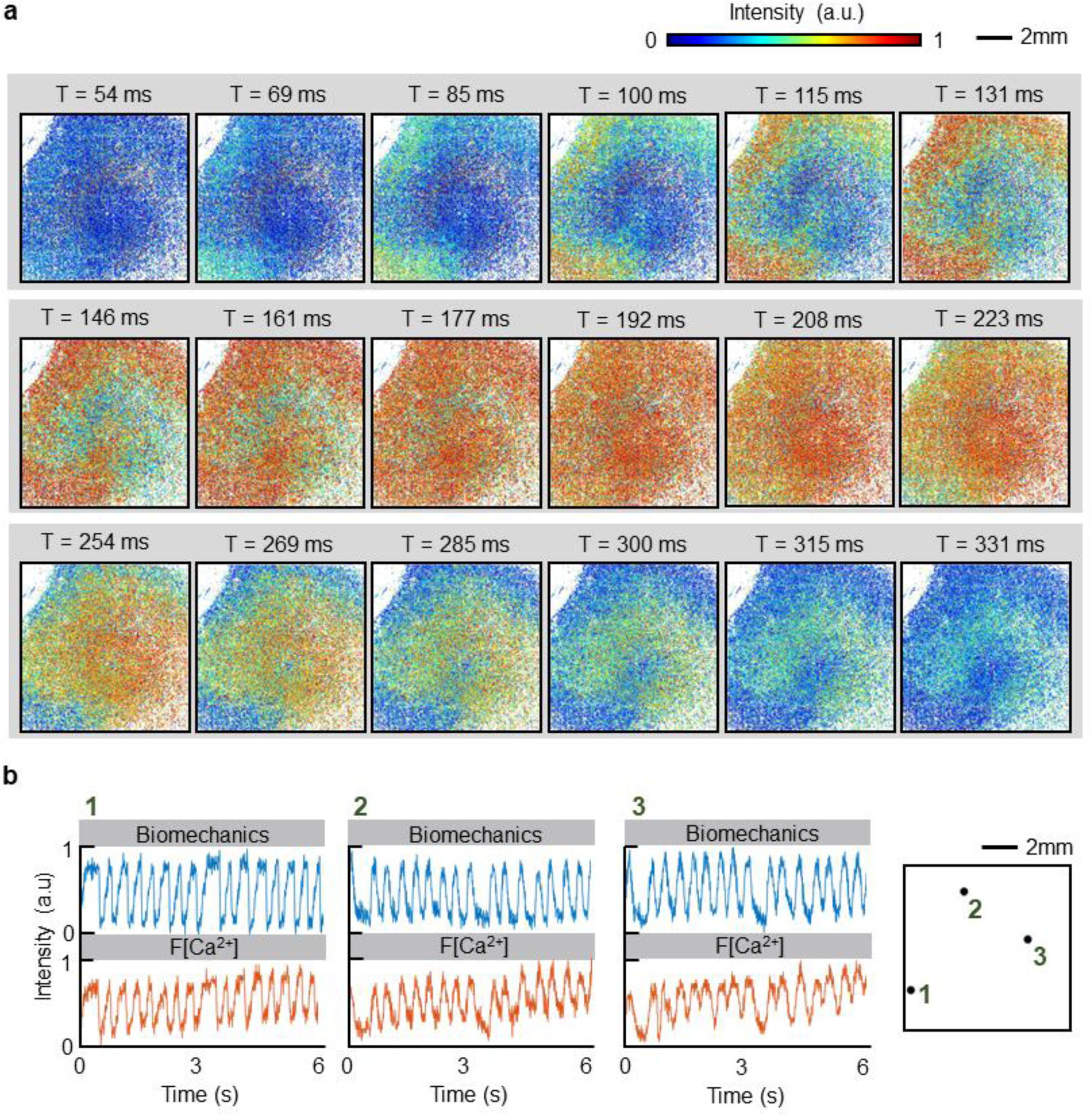
Simultaneous measurements of calcium ions concentrations and biomechanical dynamics. **a.** Time-course intensity distributions, revealing the biomechanical dynamics measured by our platform using diffractive elements. A spiral wave is clearly visible across the entire field of view. **b.** Time-course intensities indicating Ca2+ concentration (Orange curves, labelled as “F[Ca2+]) and biomechanical dynamics measured by our platform (blue curves) from various locations. It is evident that the fluorescent signal and the biomechanical signal exhibit similar profiles, including certain elongated periods.

It should be noted that with our current homemade fluorescent imaging setup, we are unable to capture changes in the fluorescent signal that reflects Ca^2+^ concentration across the entire field of view. Only a few pixels exhibit discernible changes in fluorescent intensity, and these pixels are sparsely scattered, making it difficult to recognize any spatial pattern. The primary limitations stem from the low signal nature of optical mapping using fluorescent dyes. Specifically, the signal relies on the “change” in fluorescent intensity to track change in ion concentration or transmembrane voltage. However, typically, the fractional change of these fluorescent dyes is not significant (less than 30% for most dyes)^7,24^. Moreover, the high-speed feature of dynamical cellular activities imposes an ultimate restriction on the exposure time of the image sensor (maximum exposure time = 1/frame rate), resulting in a low signal due to the short exposure time. To address these challenges, conventional optical mapping often relies on specialized equipment to collect fluorescent signals for large-area measurements during short exposure periods. For example, the cameras adopted in optical mapping often have extremely large pixel sizes (ranging from a few tens of micrometers to a few hundred micrometers per pixel) but small pixel numbers (range from 256×256 to 16×16)^26^. The large pixel size requirement also limits the spatial resolution of conventional optical mapping approaches.

Although the dynamics of Ca^2+^ concentration cannot be profiled with fine resolution over a large area in our current setup, our experiment represents the first demonstration of detailed mechanical wave propagation and the corresponding calcium ion transient. It also underscores our platform’s ability to provide high spatial resolution for recording the biomechanical dynamics. In the future, we anticipate upgrading our system to incorporate a high-quality camera specialized for optical mapping, thus enabling the acquisition of changes in fluorescent intensity across the entire field of view. This advancement will allow for simultaneous mappings of ion concentration or transmembrane voltage alongside the biomechanical dynamics measured by our platform. Consequently, this technology can provide rich information about cellular activities, serving as an invaluable tool to study the complex mechanisms underlying the interrelationships between various bio-properties.

## Discussions

In this paper, we present a novel platform for the direct measurement of large-area biomechanical dynamics. This platform harnesses the independent operation of optical diffractive elements over large areas in parallel. By eliminating the need for fiducial markers to track force-induced displacements, our approach converts force into optical properties, facilitating straightforward measurement over large areas. This is in contrast with conventional traction force microscopes, where pinpointing minimal displacements greatly restricts the field of view and small errors in displacements can lead to significant deviations in traction force recovery^11,12^ While quantitative information regarding cellular traction forces is not provided in this paper, our platform has the potential to deduce both magnitude and direction of force from measured diffraction patterns. This can be realized through the development of a comprehensive calibration model that establishes the relationship between diffraction pattern and displacements. To execute this, devices with pre-patterned displacements can be fabricated and measured, allowing for the collection of labeled data that correlates diffraction patterns with known displacements to construct a look-up calibration table. This method can provide a large amount of labelled data to train machine learning models to further enhance accuracy, albeit at the expense of laborious measurement efforts. Once the model linking the diffraction pattern and displacement is established, the relationship between displacements and traction forces can be readily determined using solid mechanical simulations^19^. Consequently, traction force can be retrieved using these two calibrated models.

On the other hand, ensuring the yield and uniformity of devices is crucial for expanding the field-of-view of the proposed platform. Currently, this is limited by the PDMS stamping processing adopted in the fabrication, which could benefit from improved pressure control and optimized PDMS buffer size. Moreover, our platform can measure other cell types, such as human-induced pluripotent stem cell-derived cardiomyocytes, for broader biological studies and discoveries. Finally, while simultaneous mapping of calcium concentrations and biomechanical cannot be achieved in our current setup, a future upgraded platform incorporating a high-quality camera specialized for optical mapping holds great potential to concurrently measure multiple bio-indicators over large area.

## Methods

### Device fabrication

The process flow of device fabrication is displayed in **Supplementary Figure S3** and described as follows. First, SU8 posts are patterned on a glass substrate using standard photolithography. These posts serve as spacers and mechanical support for subsequent PDMS stamping. Next, photoresist AZ5214 is patterned and aligned with the SU8 posts to mold the PDMS. Uncured PDMS is then poured onto the patterned substrate and covered with a polyvinyl acetate (PVA)-coated glass. PVA is a water-soluble polymer and serves as a sacrificial layer during the stamping process. A constant pressure (23 kpa) is applied on PVA-coated glass through a PDMS buffer. Then, after the PDMS is cured, the PVA-coated glass is removed by soaking the sample in deionized water. Reactive ion etching (Oxford RIE) cleans and etches the PDMS residue until photoresist AZ5214 is exposed. Subsequently, the device is soaked in acetone solution and put in the ultrasound bath for 5 seconds to strip the photoresist AZ5214. The PDMS disks, which are the key component to create displacements, are formed. Afterward, thin layers of metal films (SiO2/Ti/Al/Ti/SiO2) are deposited on the top and bottom of the PDMS disk via directional evaporation deposition. Aluminum is chosen for its high reflectivity in visible and near-infrared spectra. Finally, to ensure a flat surface, which is favored by most cells to develop focal adhesions physiologically, the PDMS stamping process is conducted again to fill PDMS to the air gaps in between PDMS disks. The final device is complete after removing the PVA-coated glass.

### Measurement Setup for Biomechanical Dynamics and Fluorescent Imaging

To keep seeded cells in their native environment for long-term, continuous monitoring of their behaviors during growth and maturation, a homemade stage top incubator is integrated into the platform. This incubator ensures a constant temperature environment and supplies the desired gas mixture optimized for cell growth. In our experiments, a constant temperature of 37°C and a humid gas mixture (95% air, 5% CO_2_) is adopted unless otherwise specified. The chamber of the stage top incubator comprises a heatable mounting frame with a 47.5mm x 21mm observation window (Zeiss, heatable mounting frame KH-L S) and a plastic cover with slidable opening and gas inlet (Zeiss, CO_2_ cover micromanipulation K). The temperature control module and the CO_2_ module are connected to a desktop and can be controlled remotely.

A deep red LED (wavelength = 660 nm) is selected for the experiments to minimize light scattering in the biological tissue by default. To offset the image plane for easy configuration of devices and the image sensor, a 90° prism mirror is placed below the stage top incubator to project the transmitted diffraction patterns to the camera lens (0.727x magnification) and the image sensor (Emergent Vision Technologies, HB25000-SB-M). Data acquisition is automated using switches and an Arduino circuit board as the control circuit. In experiments involving fluorescent images, the wavelength of LED (520-525nm) is chosen to match the excitation spectrum of the dye. A separated CMOS image sensor (IDS, UI-3080CP-M-GL, Rev.2) is utilized to record fluorescent signals. To synchronize the two cameras during data acquisition, they are fed using the same trigger signal. The dichroic mirror (Thorlabs MD568) and emission filter (Thorlabs FBH580-010) are used in accordance with the fluorescent imaging setup. **Supplementary Figure S4** illustrates the schematics and a photograph of the measurement setup described above.

### Sample preparation and cell seeding

To prepare the device for cell seeding, it undergoes several treatment steps. First the device is first rinsed with 75% isopropyl solution (IPA) followed by deionized water, and then dried with nitrogen. Subsequently, PDMS well with the desired size is bonded to the device after oxygen plasma treatment. The PDMS well sever to confine the culture medium during experiments. After bonding, the device is dried in a vacuum chamber with calcium chloride (as desiccant) overnight. Prior to seeding cells, the device surface is coated with a dopamine (Alfa Aesar Chemicals) solution (0.01%, w/v in 1M Tris-HCl buffer, Boston BioProducts, Inc.) at 4°C for an hour. Then, the device is washed twice with phosphate buffered saline (PBS) solution (Corning Inc.) and deionized water, respectively. Then, a 35-minutes ultraviolet (UV) sterilization is conducted. Once sterilized, Geltrex (83 µL/mL, Thermo Fisher Scientific Inc.) is coated on the device for 1 hour immediately before seeding cells. Neonatal rat ventricular cardiomyocytes (NRVMs) are then seeded onto the device with a density of 0.2 - 0.25 millions/cm^2^ to form a confluent monolayer tissue sheet over a 4 cm^2^ device area (active area ∼ 1cm^2^). Typically, the cells begin beating 24 to 48 hours after seeding. A culture medium containing 5% Fetal Bovine Serum (FBS) in Dulbecco’s Modified Eagle Medium (DMEM) is used for NRVMs. Fresh medium is changed every other day.

### Loading protocol of fluorescent dye

In our experiments, high-affinity calcium ion dye (Rhod-2) is utilized for fluorescent imaging. The 1mM stock solution of Rhod-2 is prepared by mixing 44.5 µL dimethyl sulfoxide (DMSO, Thermo Fisher Scientific Inc) with 50 µg Rhod-2 AM (Invitrogen). Following this, 44.5 µL Pluronic F-127 (20% w/v solution in DMSO, Thermo Fisher Scientific Inc) is added to the stock solution for assisting the dispersion of the nonpolar AM ester. For fluorescent imaging, 5 µL of the 1mM stock solution is diluted in 1mL of buffer solution (DPBS, calcium, magnesium, Gibco™, 14-040-133) and incubated with the cells seeded on the device, The final concentration of Rhod-2 AM is 5 µM. After incubating for 1 hour, the buffer solution containing the dye is removed, and the sample is rinsed with DPBS twice. The cell culture is incubated for an additional 30 minutes before fluorescent imaging is performed.

## Data analysis and statistics

Data from the image sensors was processed and analyzed using custom MATLAB codes. No statistical method was used.

## Data availability

Data associated with this study is available upon request from the authors.

## Code availability

Code associated with this study is available upon request from the authors.

## Acknowledgements

This research is supported by the National Institute of General Medical Sciences (NIGMS) of the National Institutes of Health (NIH) R01GM127985.

## Autor contribution

Y.J.L. designed and fabricated the devices, built the experimental setup, conducted experiments, developed the codes for data analysis, and wrote the manuscript; X.H.M.T. contributed to the device design, fabrication, experimental setup, biological experiments, and data analysis; Y.W. cultured and seeded the cells onto the devices. P.S.C. contributed to the development of the fabrication process, experiments, and data analysis; X.Z assisted in building the experiment setup and conducting cells experiments; T.H.W. assisted in fluorescent imaging and biological experiments; T.Y.W. assisted in the design of the experimental setup; M. A T. supervised and supported biology experiments. A. D. supervised and supported biological experiments and provided cardiomyocytes. P.Y C. conceived the device concept, supervised the project, and revised the manuscript. All authors provided input at various stages during writing.

## Competing interests

The authors declare no competing financial interests.

## Supplementary Materials

Figure S1 to S4

Movie S1 and S2

## Notes

### Competing Interest Statement

The authors have declared no competing interest.

