## Supplementary Material for "Label-Free Optical Mapping for Large-Area Biomechanical Dynamics of Multicellular Systems"

**This PDF file includes:**

1. **Diffraction patterns of diffractive elements and representative pixel extraction.**
2. **Heterogeneous behaviors in localized areas enabled by high spatial resolution of the platform.**
3. **Fabrication process flow**
4. **Measurement setup of biomechanical dynamics and fluorescent imaging**
5. **Legends for Movies S1**
6. **Legends for Movies S2**

**Other supporting materials for this manuscript include the following:**

**Movies S1 and Movies S2**

1. **Diffraction patterns of diffractive elements and data extraction**

**Figure S1a** displays the microscope image of a fabricated device. The SU8 posts form a grid surrounding the circular diffractive elements to provide uniform mechanical support during PDMS stamping. They are highlighted in orange color in one inset. Since they are usually larger and taller than the diffractive elements, their diffraction patterns are brighter than those from diffractive elements and can be easily identified and removed during post-processing. Each diffractive element will generate a diffraction pattern depending on the applied cellular traction force and will change over time according to the mechanical behaviors of cells seeded on top. The camera sensor has an 8-bit depth; therefore, the intensity reading measured in the platform will span from 0 to 255. To clearly visualize the biomechanical dynamics, we define the representative pixel of a diffractive element to be the pixel that undergoes the maximum intensity change over time. **Figure S1b** presents an example of a 3-by-3 diffraction pattern with their representative pixels marked by blue solid circles. **Figure S1b** shows the processes to extract activation time and activation duration from the representative pixels. To distinguish the signals generated by cellular activities from the fluctuated noise, the diffractive element is considered to be "activated" only when its representative pixel has intensity change larger than a certain threshold ($\Delta I > I_{thres}$), determined according to the noise levels. Then, the intensity of representative pixel is normalized to lie between zero and one. Mimicking the common definition of activation potential duration (APD), in conventional optical mapping, we define the activation duration to be the interval between the time where the intensities are half of its maximum value (Normalized intensity = 0.5). It is denoted as AD50 in **Figure S1b (ii)**. The activation duration is to characterize how long the beating lasts at a certain location On the other hand, the activation time is a figure-of-merit to characterize the timing when the beating wave arrives at a certain position, it is often defined as the point with the steepest change of fluorescent intensity and can be obtained by finding the maximum value of the first derivative. Here, to enhance the accuracy from our discrete data points, the rising part of the normalized intensity is fitted by a logistic curve and the midpoint of the logistic function (**$t_0$**) is considered as the activation time (**Figure S1b (iii)**).

The representative pixels discussed above allow us to quickly retrieve some crucial properties of biomechanical dynamics including beating pattern, activation duration, activation time, and dominant frequency. Alternatively, the diffraction pattern, along with the intensity sum of all the pixels in a diffraction pattern, contains rich information for extracting both the magnitude and direction of cellular traction force quantitively. In the future, the development of a comprehensive calibration model that establishes the relationship between diffraction pattern and displacements can provide more quantitative analysis basis of cellular traction forces.

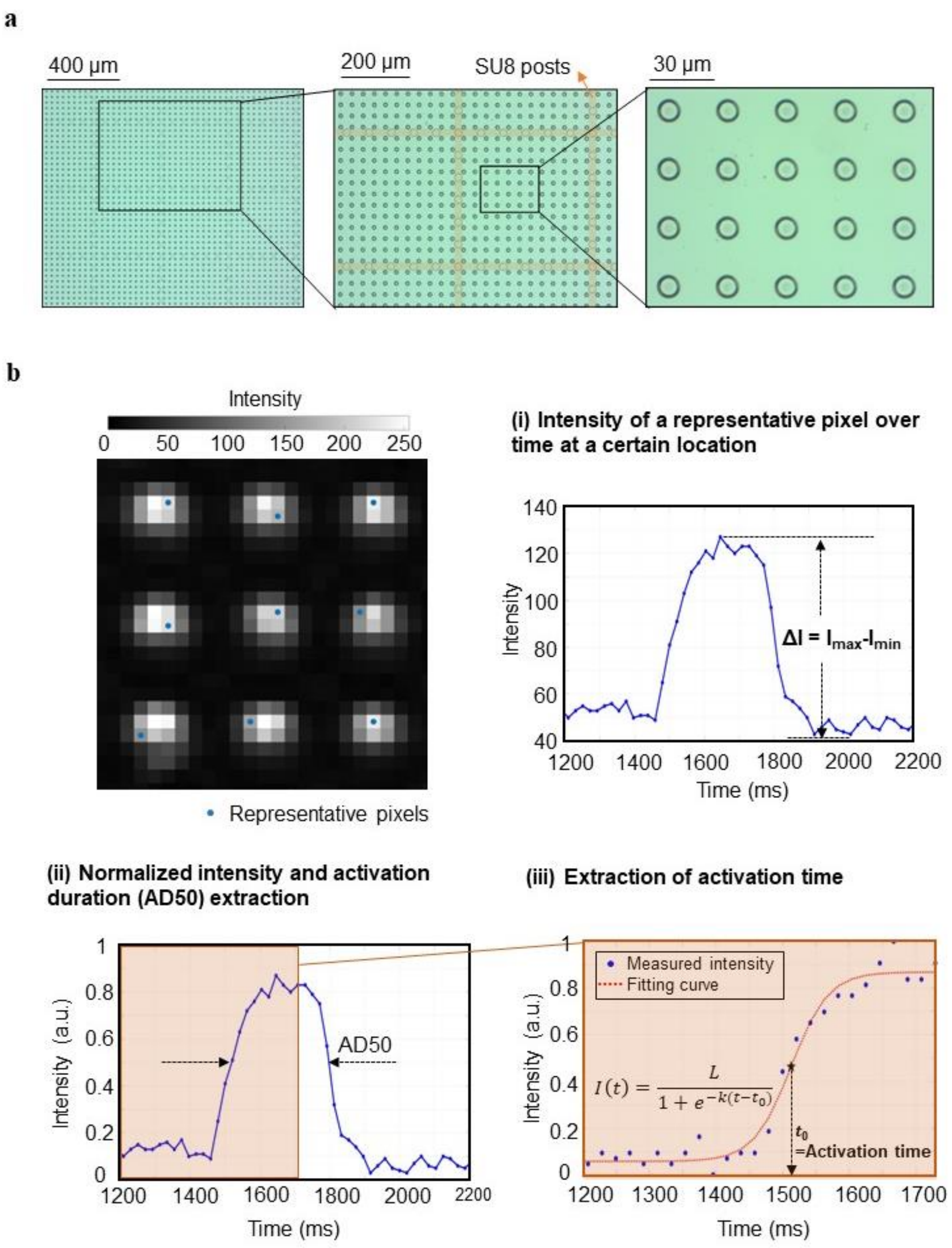


**Supplementary Figure S1. Diffraction patterns of diffractive elements and data extraction. a.** Microscope images of device. **b.** Representative pixels identified for 3-by-3 diffraction patterns, along with the extraction processes for activation duration and activation time: (i) Intensity of a representative pixel over time at a specific location (ii) Normalized intensity and extraction of activation duration (AD50) (iii) Extraction of activation time.

### 2. Heterogeneous behaviors in localized areas enabled by high spatial resolution of the platform.

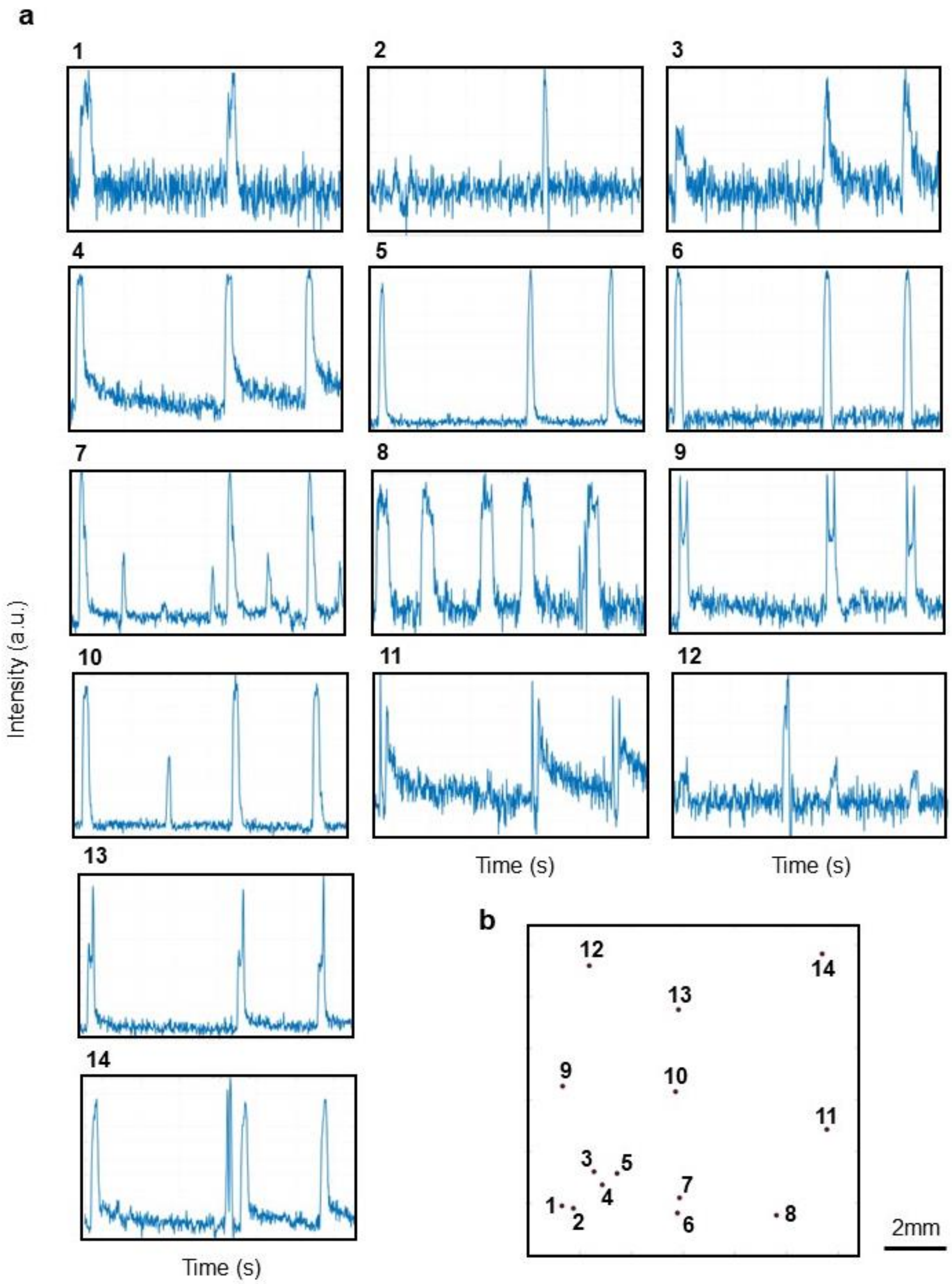


**Supplementary Figure S2. Heterogeneous behaviors in localized areas enabled by high spatial resolution of the platform. a.** Selected heterogeneous traces depicting the intensity change over time at various locations for the three consecutive cardiac beatings presented in **Fig. 2**. Thanks to the high spatial resolution of our platform, local heterogeneities can be identified with unprecedent details while recording global beating patterns. This is a powerful tool to investigate collective behaviors and track individual cellular activities simultaneously. The x-axis represents time with a span of 15 seconds while the y-axis represents normalized intensity ranging from 0 to 1. **b.** The corresponding locations of each point shown in **a.** The entire field-of-view is 10.6mm by 10.6mm.

#### 3. Process flow of device fabrication

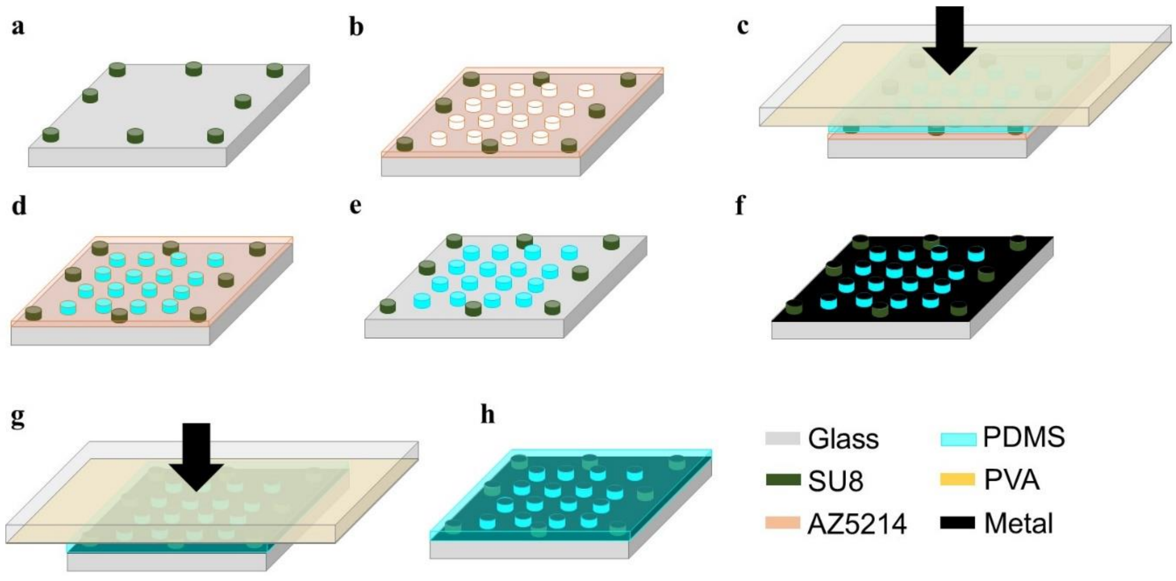


**Supplementary Figure S3. Process flow of device fabrication. a** Patterning SU8 posts as spacers and mechanical support for subsequent PDMS stamping. **b.** Patterning photoresist AZ5214 and aligning it with the SU8 posts to mold the PDMS. **c.** Stamping PDMS: Applying a constant pressure (23 kpa) on PVA-coated glass through a PDMS buffer. PVA is water-soluble and serves as a sacrificial layer during the stamping process. **d**. Removing PDMS residues via dry etching. **e.** Stripping photoresist AZ5214 using acetone. The PDMS disks, the key component for creating displacements, are formed. **f**. Isotropic metal deposition. **g.** Stamping PDMS: filling the air gaps between PDMS disks with PDMS. A second stamping ensures a flat surface, favorable for physiological development of focal adhesions. **h.** Final device. PDMS: polydimethylsiloxane. PVA: polyvinyl acetate.

### 4. Measurement setup of biomechanical dynamics and fluorescent imaging

a

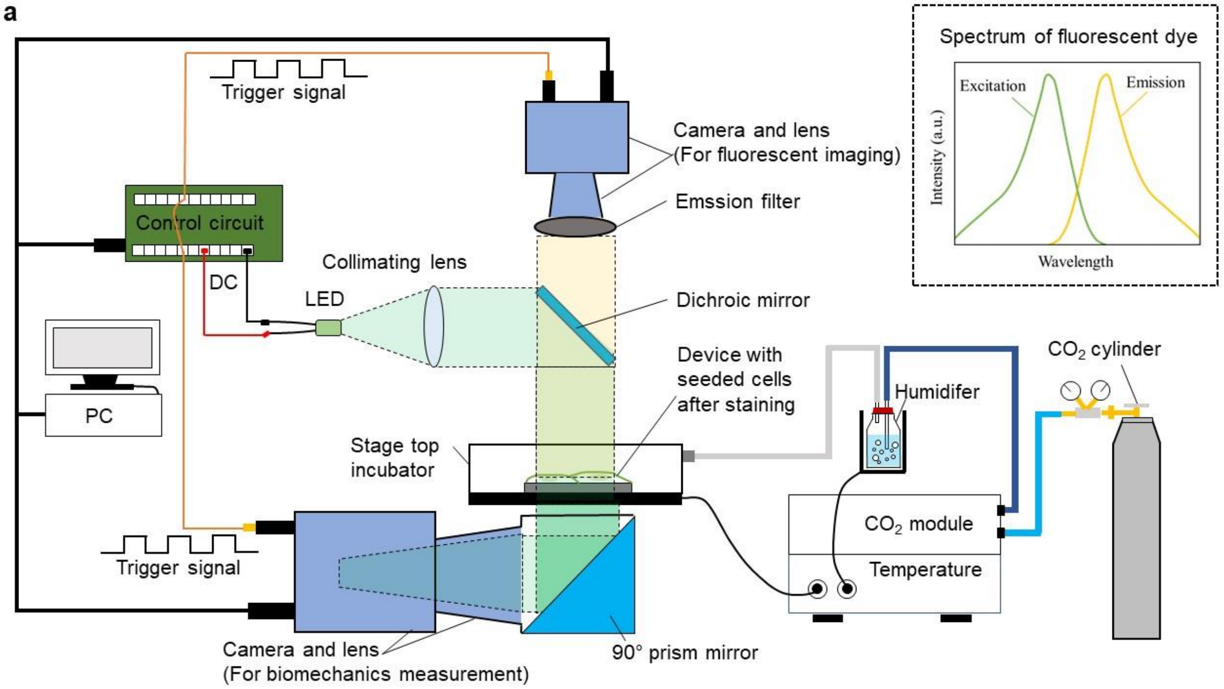


b

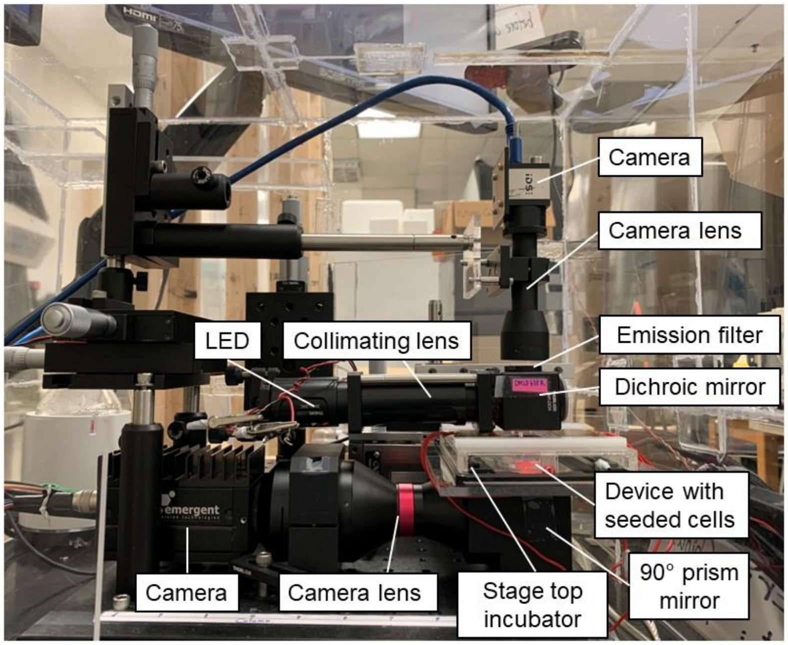


**Supplementary Figure S4. Measurement setup for biomechanical dynamics and fluorescent imaging.** **a.** Schematic of the measurement setup: A LED is collimated to illuminate cells seeded on top of the device. Transmitted diffraction patterns are directed to the camera below via a 90° prism mirror. For fluorescent imaging, a separated image sensor is utilized to record fluorescent signals in the reflected direction. To synchronize the two cameras during data acquisition, they are fed using the same trigger signal. Data acquisition is automated using switches and an Arduino circuit board. The dichroic mirror and emission filter are incorporated for fluorescent imaging. A homemade stage top incubator is integrated to the setup to ensure a constant temperature environment and supplies the desired gas mixture optimized for cell growth **b**. Photo of the measurement setup.

### 5. Legends for Movies S1

**Movie S1** (as a separated file) **Real-time visualization of transient mechanical wave propagation over large area with high spatial resolution**. A sub-million number of neonatal rat ventricular myocytes (NRVMs) were seeded onto the devices, and their cardiac activities were monitored. The video illustrates two consecutive cardiac beats: the first beat is shown on the left-hand side (with a timeframe from 1.08 to 3.13 seconds) while the second beat is presented on the right-hand side (with a timeframe from 8.79 to 10.83 seconds). The video plays at a rate of 24 frames per second, which is half the recording rate of 48 frames per second). The color representing the normalized intensity over time extracted from each activated diffractive element. The field-of-view is 10.6mm by 10.6mm. This video highlights the rapid changes in mechanical wave patterns between beats across a large area, demonstrating the demand for large-area measurement and the capabilities of our platform.

### 6. Legends for Movies S2

**Movie S2** (as a separated file) **Continuous monitoring of mechanical dynamics during cardioversion via electrical stimulation**. A sub-million number of neonatal rat ventricular myocytes (NRVMs) were seeded onto the devices. The video illustrates the cardiac activities from the same sample before and after electrical stimulation, displayed on the left-hand side and right-hand side, respectively. The video plays at a rate of 24 frames per second, which is half the recording rate of 48 frames per second). The color representing the normalized intensity over time extracted from each activated diffractive element. The field-of-view is 10.6mm by 10.6mm.
