## Supplementary figures and images for "Label-Free Optical Mapping for Large-Area Biomechanical Dynamics of Multicellular Systems"

### VideoS2

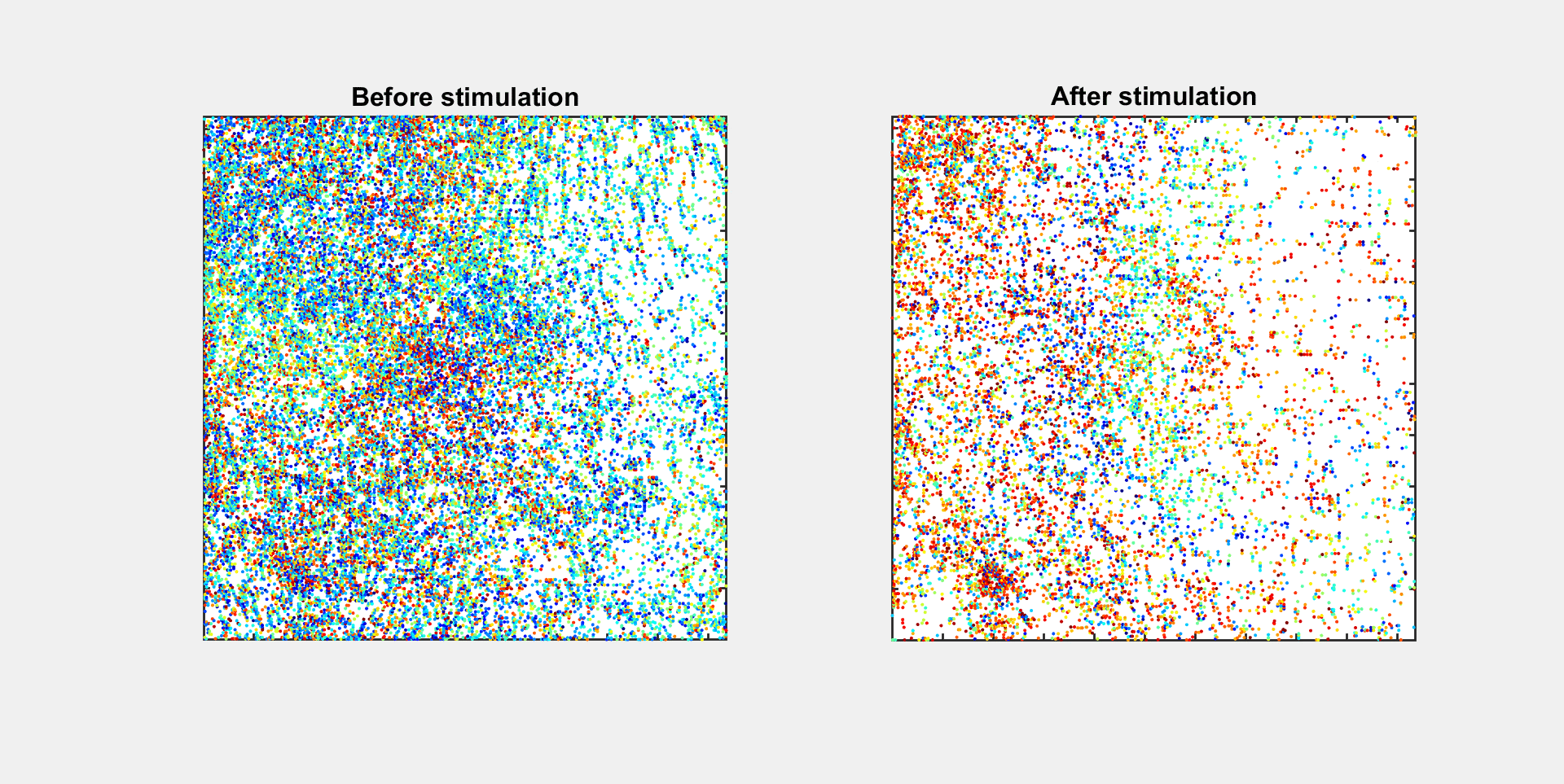
